# Cold Atmospheric Plasma for Alzheimer’s disease

**DOI:** 10.1101/2025.11.07.687156

**Authors:** Xiao Chen, Zhuonan Yu, Chunyan Du, Yueye Huang, Zhaowei Chen, Zhen Xu, Zhitong Chen

## Abstract

Lightning has been proposed as a pivotal energy source driving prebiotic chemical evolution and early life processes. Similarly, external physical stimuli such as sound and light have been shown to modulate neurophysiological activity and mitigate Alzheimer’s disease (AD) pathology by promoting amyloid-β (Aβ) clearance. Here, we demonstrate that cold atmospheric plasma (CAP) acts as a joule-level analog of lightning, integrating light, sound, and reactive species to modulate AD progression in a mouse model. Using multimodal analyses, we show that CAP treatment enhances key neuroimmune signaling pathways in an AD murine model, including microglial activation, without inducing pathological alterations at the functional, transcriptomic, or proteomic levels in healthy mice. These findings highlight CAP as a safe and efficacious modality for modulating neurodegenerative processes, establishing a foundation for its therapeutic translation in AD and related disorders.

## Introduction

Alzheimer’s disease (AD) is a progressive neurodegenerative disease characterized by the accumulation of amyloid beta (Aβ) plaques and the entanglement of hyperphosphorylated tau protein^1^ ^2^. These pathological changes collectively lead to synaptic dysfunction, neuropathy, and cognitive decline^3^. Globally, AD affects millions of people and is a major challenge in the field of public health^4^. Although decades of research have explored targeted therapeutic strategies for amyloid and tau proteins^5^ ^6^, these methods have limited effectiveness in alleviating symptoms or delaying disease progression, partly due to the incomplete understanding of the complex multifactorial pathological mechanisms of AD^3,7^. Therapies targeting amyloid proteins have not efficiently improved patient prognosis, suggesting that a more comprehensive strategy is needed for AD treatment.

Currently, there is no known cure for AD, and the efficacy of existing drug treatments is limited, highlighting the urgent need for new non-invasive therapies. Among emerging therapies, non-invasive physical intervention strategies have attracted growing attention. In recent years, studies have shown that there is impaired gamma frequency oscillation (30-100 Hz) in AD patients and mouse models, which is closely related to cognitive dysfunction ^8^. Gamma frequency sensory stimulation has shown potential as an innovative non-pharmacological intervention, inducing brain gamma activity through external stimuli such as 40 Hz light flashes or auditory pulses to alleviate AD pathology ^9^ ^10^. In experimental models such as 5xFAD or APP/PS1 mice, 40 Hz light or sound stimulation not only reduces Aβ burden, but also improves behavioral test performance, including cognitive function and motivation recovery ^11^.

Lightning is a common atmospheric phenomenon in nature, which is essentially a plasma discharge^12^. As an active participant in the origin of life, lightning produces light, sound, and reactive nitrogen - and oxygen-containing substances^13^ ^14^, which play a role in nitrogen fixation^15^, carbon dioxide reduction ^16^, and the production of life precursor molecules such as ammonia (NH^4^), nitrate, and formic acid ^12^ ^17^. Despite its biological relevance, the potential of lightning-simulating plasma phenomena in therapeutic applications remains less explored.

Here, we propose cold atmospheric plasma (CAP) as a controllable, joule-level analog of lightning that integrates photonic, acoustic, and reactive chemical components as a multimodal physical stimulus.^16–18^ Our findings demonstrate that CAP effectively modulates key signaling pathways implicated in AD progression, including microglial activation, while exhibiting no pathological alterations in healthy mice at the functional, transcriptomic, or proteomic levels. CAP does not induce inflammation or oxidative stress, highlighting its specificity, biocompatibility, and non-invasive safety profile. Collectively, these results establish CAP as a promising and safe physical intervention for neurodegenerative diseases, offering a novel paradigm for modulating complex brain pathophysiology.

## Results

### The CAP-derived reactive species composition is different between different sources

Our plasma stimulation system was designed to deliver not only light and sound cues to mice but also gaseous plasma components. To achieve this, an air pump was integrated to introduce plasma effluents into the chamber, while an open outlet at the upper end maintained air circulation (**Figure 1A**). Different carrier gases yield distinct plasma chemistries, generating diverse reactive species that influence both plasma behavior and potential biological effects. As expected, air-based plasma produced abundant reactive oxygen and nitrogen species (RONS), including characteristic emission lines of O I, OH, N_₂_, and N_₂_^⁺^ (**Figure S1A**). Excited atomic oxygen (O I) can further react with molecular oxygen to form ozone, which at high concentrations may be harmful to the respiratory system. Helium-based plasma primarily generated reactive nitrogen and oxygen species with prominent helium-related spectral lines (**Figure S1B**). Nitrogen plasma yielded a higher proportion of reactive nitrogen species (RNS) (**Figure S1C**), whereas argon plasma produced both RNS and ROS together with argon-specific emissions (**Figure S1D**). To best replicate natural atmospheric conditions, air plasma was selected for subsequent experiments.

**Figure 1.**
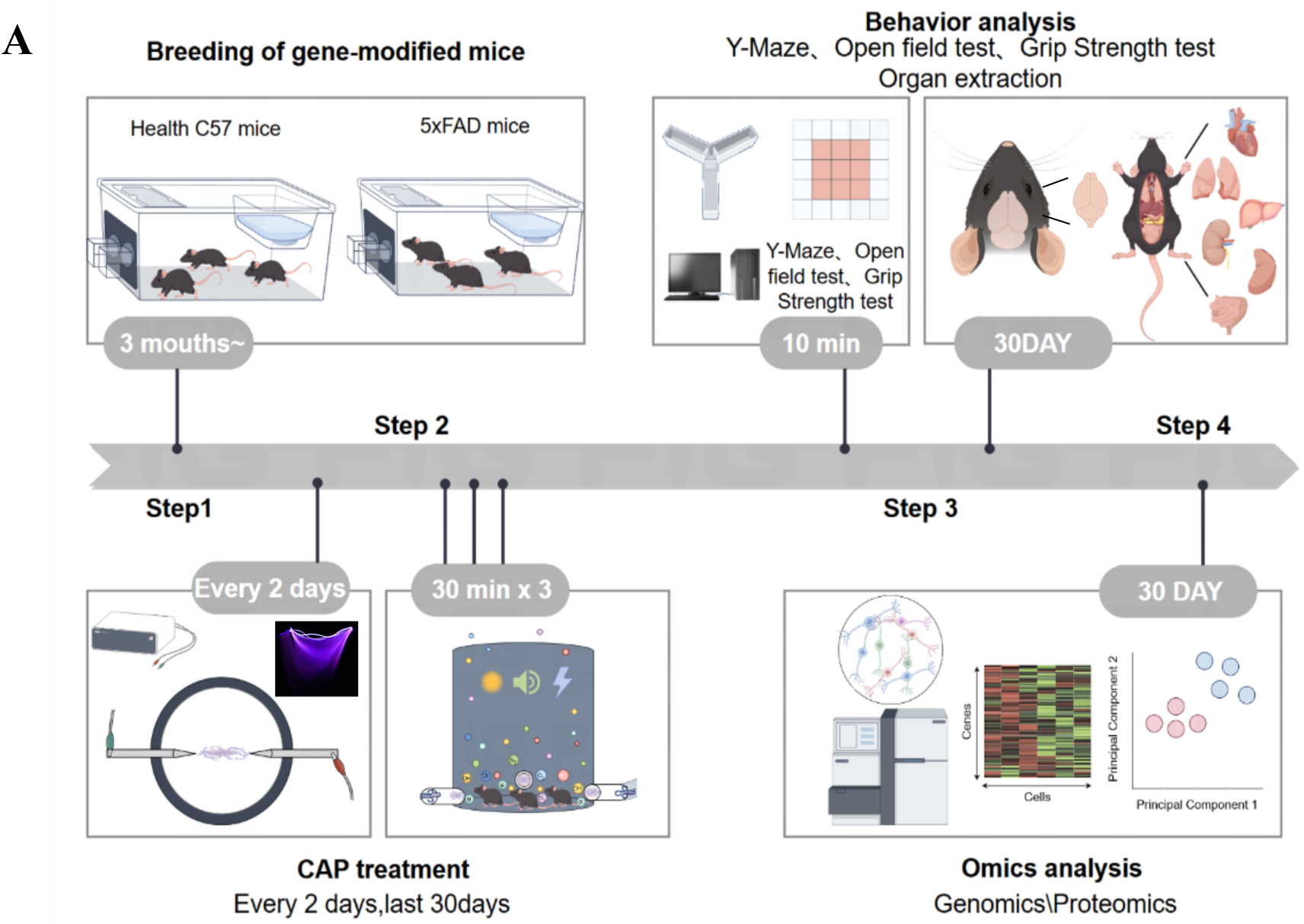
Schematic and characteristics of the CAP device. Mice were acclimated to the environment and then randomly assigned to either a control group (no CAP exposure) or a treatment group receiving CAP for 30 minutes every other day over one month in dedicated devices. At the conclusion of the treatment period, behavioral tests were conducted, and brains and major organs were collected for pathological staining and histological analysis to comprehensively evaluate the treatment effect and safety.

### CAP activates disease-associated signaling pathways in an AD mouse model

One of the less explored frontiers for the clinical application of CAP is the neurodegenerative disease. We employed the 5xFAD mouse strain, a well-established model of AD, to investigate the molecular effects of CAP stimulation. Mice were randomly allocated to the control group or CAP treatment group, acclimated to the experimental environment, and underwent CAP intervention for 30 minutes every other day in a dedicated CAP device over a one-month period. RNA-sequencing of whole-brain tissues from CAP-treated and control 5xFAD mice identified 16006 genes, among which 261 genes were differentially expressed, distinctly segregating samples into two groups based on CAP treatment (**Figure 2A**). Several representative upregulated genes were validated in the brains of CAP-treated AD mice, such as components of the complement system (*C1qa*, *C1qb*, *C1qc, C4b*), the phagocytosis system (*Fcgr1 and Cd68*), and the Tyrobp causal network (*Trem2 and Tyrobp*) (**Figure 2B**). As these pathways are closely associated with microglia activation and phagocytosis, we postulated that CAP treatment might enhance the activity of microglia. Further gene ontology analysis revealed an enrichment not only in pathways associated with injury/wound healing, cytokines/chemokines, but notably in microglia-related phagocytosis and Tyr pathways within the CAP-treated group (**Figure 2C**). These results indicate that CAP exerts diverse effects on microglial gene expression, particularly in pathways relevant to phagocytosis, which is significantly implicated in Alzheimer’s disease pathology.

**Figure 2.**
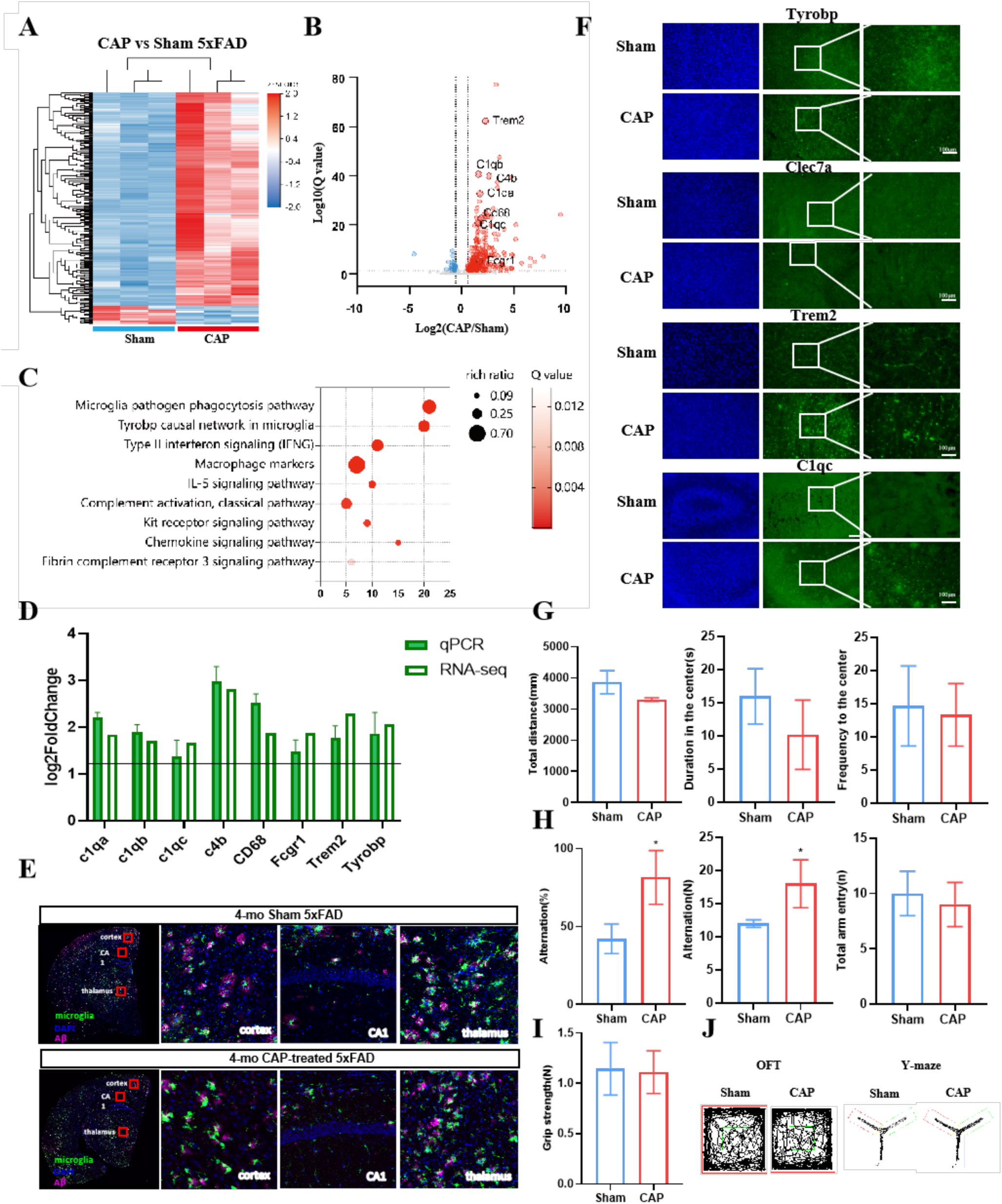
CAP activates disease-associated signaling pathways in a mouse model of AD. (A) Gene Heatmap of differentially expressed genes; red and blue represent high and low expression levels, respectively. (B) Volcano plot showing upregulated (red) and downregulated (green) genes. (C) KEGG pathway enrichment analysis of differentially expressed genes. (D) Expression levels of *C1qa, C1qb, C1qc, C4b, Fcgr1, CD68, Trem2 and Tyrobp*. (E) Immunofluorescence staining of *C1qc, Clec7a, Tyrobp,* and *Trem2* in CAP-treated and control 5xFAD mouse brains. (F) Double immunofluorescence staining of Aβ (green) and microglia (pink) in the cortex, CA1, and thalamus. (G) Open-field test results, including total distance, duration in the center, and frequency to the center. (H) Y-maze test results, including alternation (%), alternation number, total arm entry number, and (I) Grip-strength test results. (J) Behavioral experiment trajectory diagram. Data are presented as mean ± SE (n = 3 mice per group). Statistical significance was calculated via Student’s *t*-test. \**P* < 0.05.

Next, to validate transcriptomic findings, **qPCR** confirmed increased expression of complement, phagocytic, and Tyrobp-network genes following CAP exposure (**Figure 2D**). Consistently, immunofluorescence staining for *C1qc*, *Cxcl7a*, *Tyrobp*, and *Trem2* demonstrated elevated protein levels in CAP-treated mice (**Figure 2E**), supporting microglial activation. We performed fluorescence staining for Aβ protein and microglia. In the hippocampal region (CA1), green fluorescence increased and pink fluorescence decreased, suggesting microglia activation and Aβ protein reduction (**Figure 2F**). These results indicate that CAP facilitates Aβ clearance via microglia-mediated phagocytosis.

Given the transcriptomic and proteomic alterations observed in CAP-treated AD brains, we next examined whether these molecular changes translated into behavioral effects. Open field test, Y-maze test, and grip-strength tests were performed on 5xFAD mice treated with CAP and without CAP exposure to evaluate locomotor activity, anxiety, cognitive performance, and limb muscle strength. In the open-field test, CAP-treated mice showed no significant difference in total distance traveled compared with controls, indicating that CAP treatment did not impair locomotor function. Similarly, the time spent and the number of entries in the central zone were comparable between groups, suggesting that CAP treatment would not cause anxiety-like behavior (**Figure 2G**). Because excessive RONS have been implicated in neurodegenerative pathology, we assessed cognitive outcomes using the Y-maze test. CAP-treated mice exhibited significantly higher spontaneous alternation counts and rates than those of the sham group (*P*<0.05), indicating improved spatial working memory (**Figure 2H**). Finally, the grip-strength test results revealed no significant difference between groups, confirming that CAP treatment did not damage muscular function (**Figure 2I**). Collectively, these findings demonstrate that CAP treatment does not impair motor or anxiety-related behaviors in AD mice, but instead partially enhances memory performance, consistent with its neuroprotective molecular effects.

### CAP does not induce a pathological response in healthy mice

A major concern regarding CAP application is the potential for reactive species to induce oxidative stress and inflammation, especially along the infusion route, in our case the respiratory system. To assess the biosafety of CAP, we evaluated hematological, biochemical, histological, and behavioral parameters in healthy mice exposed to CAP. Hematological analysis revealed no significant alterations in white blood cell counts or in liver and kidney function compared with untreated controls (**Figure 3A-H**), indicating the absence of systemic inflammation or organ toxicity. Histological examination using hematoxylin and eosin (H&E) staining showed no morphological abnormalities in major organs following CAP exposure (**Figure 3I)**. There were no obvious fluctuations in the body weight through the experiment (**Figure 3J)**. To further determine whether CAP affected physiological or neurological function, we performed open-field, Y-maze, and grip-strength tests in healthy mice. No significant differences were observed between CAP-treated and control groups across locomotor, anxiety-related, memory, or muscle-strength parameters (**Figure 3K-S)**. Together, these findings demonstrate that CAP exposure does not elicit pathological, inflammatory, or behavioral abnormalities in healthy mice, providing preliminary evidence of its safety and biocompatibility for potential therapeutic applications.

**Figure 3.**
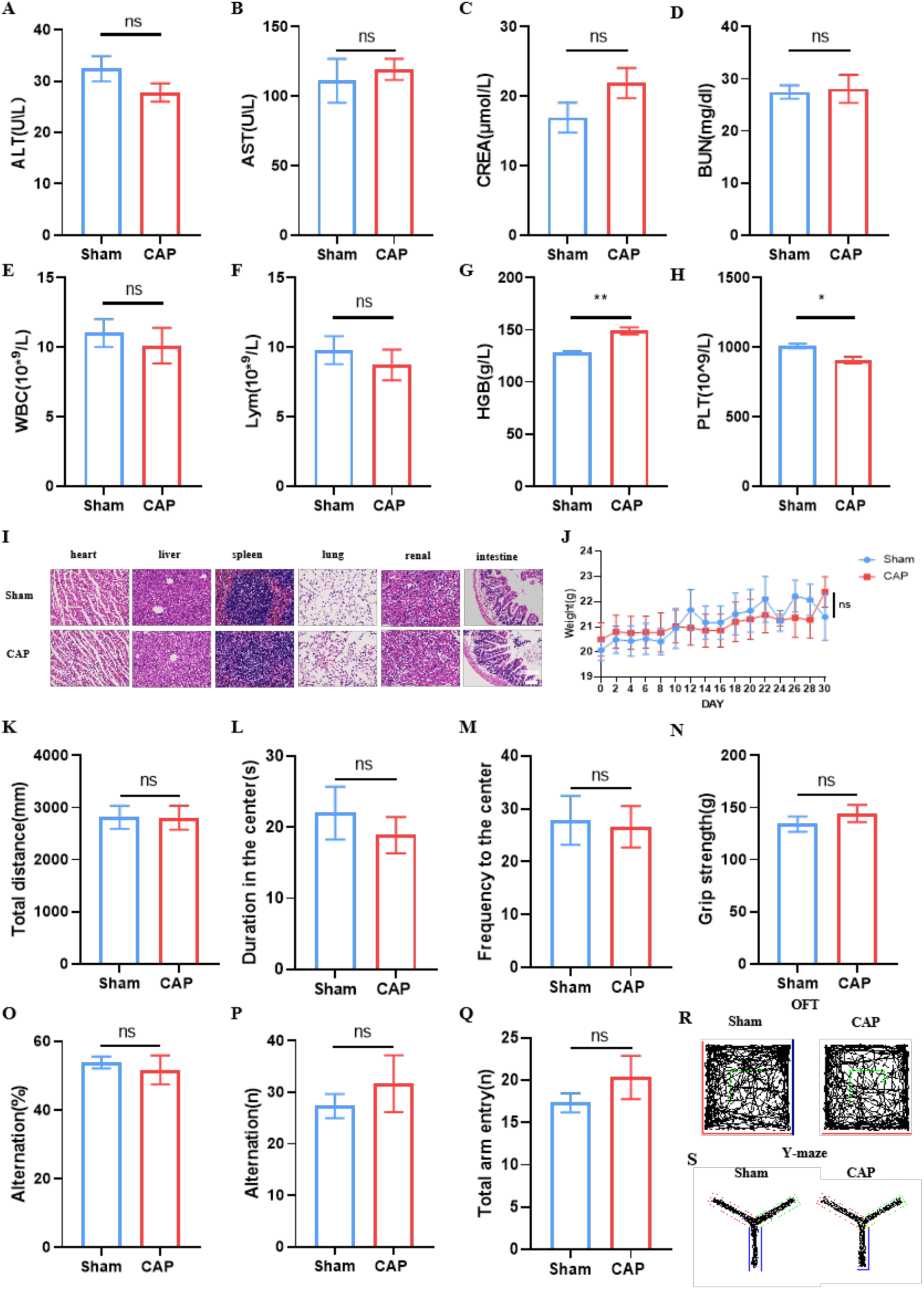
CAP does not induce a pathological response in healthy mice. (A-H) Behavioral test results for CAP-treated and sham groups. (I) H&E staining images of major organs, including the heart, liver, spleen, lung, kidney, and intestine. Scale bar, 200 μm. (J) The average body weight of mice during the experimental period. (K-Q) Blood biochemical parameters related to liver and kidney function. (R) Trajectory diagram of the open field test. (S) Trajectory diagram of the Y-maze test. Data are presented as mean ± SE (n = 3 mice per group). Statistical significance was calculated via Student’s *t*-test. \**P* < 0.05.

### CAP does not induce molecule-scale pathological responses in healthy mice

Though CAP-derived reactive species may not cause pathological alterations at the macroscopic level, they could potentially trigger subtle molecular changes. To determine whether CAP-derived reactive species induce transcriptomic abnormalities in healthy mice, we extracted RNA from multiple organs of mice treated with CAP, performed RNA sequencing, and analyzed the transcriptomic profiles. We found that inhalation of CAP-derived reactive species caused minimal transcriptomic alterations across all examined organs (**Figure 4**).

**Figure 4.**
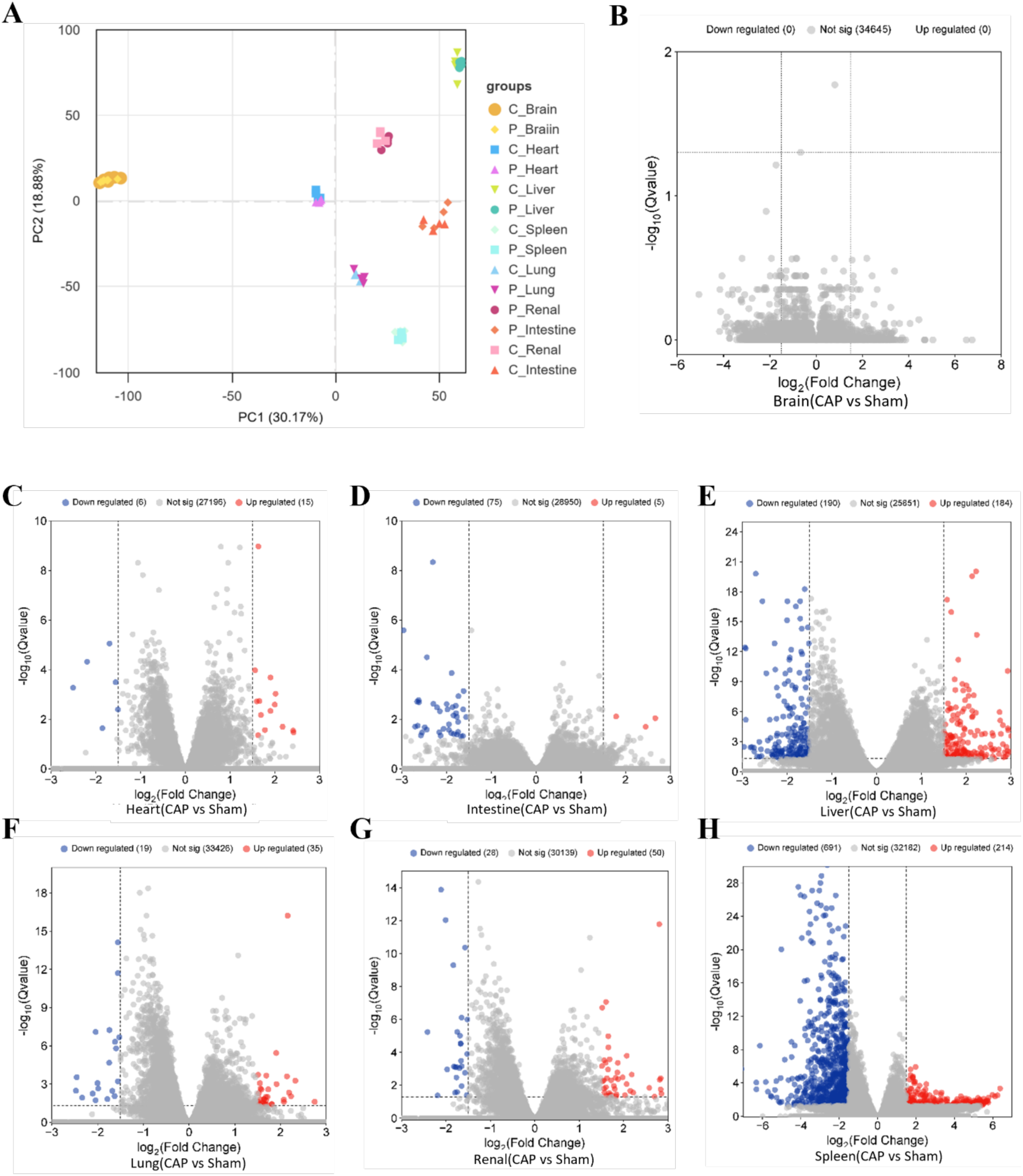
Transcriptomic analysis of CAP-treated healthy mice. (A) Transcriptomes from different organs were clustered correctly by principal analysis. (B-H) Volcanic diagram of gene expression changes. The genes that were significantly upregulated (red) and downregulated (blue) were highlighted. The dashed line represents the threshold for significance (*P=0.05*) and fold change (|log2FC|=1.5).

To further corroborate these findings, we performed proteomic analyses of brain tissues from CAP-treated and control mice. Similarly, inhalation of CAP-derived reactive species hardly resulted in negligible proteomic changes in all the organs we studied (**Figure S2**). To capture the subtle difference between CAP-treated mice and control mice, we also performed gene set enrichment analysis (GSEA) based on proteomic data, which also revealed no significant changes (**Figure S2**). Collectively, these results demonstrate that CAP-derived reactive species do not induce pathological responses at the proteomic or proteomic level, confirming the biosafety of CAP treatment.

## Discussion

Our study provides the first evidence that CAP, a novel multimodal physical stimulation, exerts therapeutic effects in an AD mouse model by modulating key neuroimmune signaling pathways, most notably microglial activation, without inducing adverse effects in healthy subjects. Microglia have been clearly identified as central coordinators of AD neuroinflammation and regulators of Aβ dynamics^18^. In AD, microglia display substantial heterogeneity. Although moderately activated microglia facilitate Aβ clearance, excessive activation can trigger chronic and harmful inflammatory reactions, exacerbating neuronal damage and synaptic loss^19^ ^20^ ^21^. The moderate activation of microglia observed following CAP exposure may therefore underlie the reduced pro-inflammatory cytokine levels and enhanced Aβ phagocytic activity in our model, effectively interrupting the inflammatory-Aβ feedback loop. This phenotype shift likely involves altered expression of key microglial receptors such as TREM2, which is closely related to Aβ recognition and clearance and microglial function in AD^22^.

The compelling safety profile of CAP represents a critical translational advantage ^23^ ^24^, aligning with the established tolerability of non-invasive neuromodulation techniques, including transcranial pulse stimulation (TPS) and gamma stimulation. Clinical trials of TPS in AD patients have consistently demonstrated cognitive and neuropsychiatric improvements with minimal adverse events, underscoring the clinical feasibility of non-invasive stimulation paradigms^25^. Such a favorable safety characteristic is paramount for clinical adoption, especially given the vulnerability of the AD population. Similarly, in our studies, CAP exposure did not induce systemic inflammation, oxidative stress, or significant transcriptomic or proteomic perturbations in healthy animals, highlighting its intrinsic biocompatibility. This aligns with the lack of serious adverse effects reported in human trials employing non-invasive gamma sensory stimulation and transcranial pulse stimulation. Consequently, CAP leverages the inherent safety benefits of non-invasive physical energy delivery, providing a strong foundation for its clinical advancement as a well-tolerated and scalable therapeutic strategy for AD, suitable for extended or repeated application regimes, required for effective disease modification.

Mechanistically, CAP represents a unique multimodal platform that integrates optical, acoustic, and plasma components into a single bioactive stimulus^26,27,28^. Based on our observed stimulus, we hypothesize that CAP-generated RONS act as primary biochemical mediators, initiating signaling cascades that modulate microglial activation and neuroinflammatory balance. RONS are known to regulate key transcriptional regulators ^29^ ^30^, such as NF-κB and Nrf2, which orchestrate inflammatory and antioxidant responses. Crucially, co-delivery of light and acoustic cues at specific frequencies may synergistically enhance the cellular responsiveness to these signaling pathways, thereby refining the accuracy of microglial regulation^31–33^. For example, previous studies have shown that pulsed light stimulation promotes beneficial microglial responses (such as increased Aβ co-localization, M1 to M2 transition^34^) and vascular Aβ clearance, while electromagnetic or acoustic pulse therapies can similarly attenuate neuroinflammation and Aβ pathology^35^. CAP thus advances these modalities by integrating multiple synergistic physical and chemical cues within a unified therapeutic framework^36^ ^37^. We propose that the concurrent exposure to photonic and acoustic energy, together with CAP-derived biologically active RONS, creates a distinctive bioelectrochemical microenvironment. Light and sound at specific frequencies may activate cells, enhance membrane permeability, or resonate with specific cell structures or signal molecules, thus amplifying the efficacy and specificity of RONS-mediated signaling on microglial targets compared with any single mode alone ^38^. Such coordinated stimulation could facilitate microglial phenotypic reprogramming^39^, enhance TREM2-dependent Aβ phagocytosis^40^ ^41^ ^22^ and lysosomal degradation, and promote more effective plaque clearance and attenuation of neurotoxic inflammation than single-mode physical interventions.

Future work should systematically dissect the molecular mechanisms and synergistic interplay among CAP’s constituent components, with particular attention to how plasma-derived RONS spectra interact with light- and sound-activated signaling networks in microglia. Long-term safety studies are also warranted to assess the cumulative effects of chronic or repeated CAP exposure. Furthermore, expanding this approach to human cellular and organoid models will be essential to validate translatability and therapeutic precision in clinical contexts.

## Conclusion

This study provides compelling evidence that CAP, a multi-modal therapeutic modality integrating optical, acoustic, and reactive chemical stimuli, exerts profound regulatory effects on AD pathophysiology in murine models. By modulating microglial signaling and inflammatory balance, CAP effectively mitigates AD-related neuropathology. Notably, comprehensive behavioral, transcriptomic, and proteomic analyses revealed no adverse alterations in healthy mice, underscoring the safety profile of CAP. Collectively, these findings establish CAP as a promising, non-invasive, and mechanistically versatile neuromodulatory strategy for neurodegenerative disease intervention, paving the way for its further optimization and clinical development.

## Method

### Animal

Animals were provided food and water by the experimental animal center of Shenzhen Advanced Technology Research Institute, Chinese Academy of Sciences (SIAT) in accordance with rodent feeding standards. Methods Animals Procedures were approved by the Research Institutional Animal Care and Use Committee and adhered to standards set forth by the Guide for the Care and Use of Laboratory Animals. All of the animal holding rooms were maintained within temperature (18–26 °C) and humidity ranges (30– 70%) described in the ILAR Guide for the Care and Use of Laboratory Animals (1996). The 5xFAD mice and C57BL/6 J mice used in the experiment were raised in a standard feeding environment, with a standard light-dark cycle of 12 hours to 12 hours (lights on at 07:00; all experiments were conducted within the light cycle).

### CAP equipment

The plasma device employed in this study is composed of two metal electrodes (1 mm in diameter), facing each other at a distance of 5 mm, to generate plasma between the pins through an applied electric field. (Figure 1A). Thermal images of the CAP device were taken using a handheld thermography camera (HIKMICRO, HM-TPH21Pro-3AQF). The discharge voltage was measured by an oscilloscope (Tektronix, TDS2024C) with a high-voltage probe (Tronovo, TR9340A). The optical emission spectrometry (OES) was characterized by a high-resolution UV/visible spectrometer (Brolight, BIM-6602A series), setting up the optical probe placed 1 cm away from the discharge area.

### CAP stimulation

The experimental flow chart is shown in Figure 1A. The mice were removed from the animal breeding room and kept in a quiet room. After getting used to the room for 1 hour, place individual mice in separate chambers. This chamber is cylindrical with circular holes with a diameter of 1cm around the bottom for placing plasma devices. The device is connected to a power source and an air pump. While the air pump blows CAP substance into the chamber, the mice also receive the light and sound generated by CAP. The top of the device is not sealed to ensure smooth airflow. The mice were divided into two groups: one control group, which did not receive CAP treatment, but were placed in the same-sized device for the same time treatment; The other group is the treatment group, which received CAP treatment and is treated once every other day for 30 minutes each time for one month.

### Tissue collection and processing for immunohistochemistry

After CAP stimulation, mice were given a lethal dose of anesthetic (isoflurane overdose), followed by cardiac perfusion with PBS (pH 7.4). Then there is PBS containing 4% paraformaldehyde (PFA). After perfusion, the skull cap was removed and the brain was stored in 4% PFA for 18-24 hours, washed in PBS, and then dehydrated in 30% sucrose until the brain tissue sank, embedded in OCT (Tissue Tek), frozen at −80 °C, and then cut at 40 μm in a low-temperature thermostat and mounted on SuperFrost glass slides. Then, perform immunohistochemistry on the tissue sections mounted on the glass slide.

### Immunohistochemistry

Perform immunohistochemical treatment on coronal brain slices using the following protocol. Firstly, wash the tissue with PBS for 10 minutes, permeabilize with 0.1% Triton X-100 in PBS for 10 minutes, block at room temperature (5% normal donkey serum and 0.1% Triton X-100 in PBS) for 2 hours, and stain overnight with primary antibody immunostaining in the blocking solution. After washing 3 times with blocking buffer for 5 minutes, add the secondary antibody to the blocking buffer at room temperature for 2 hours, and then wash 5 times with PBS for 5 minutes each time. Seal with Prolong Gold sealing agent (Thermo Fisher Scientific, P36930).

### Behavioral Assessments

To evaluate the effects of cold atmospheric plasma treatment on locomotor activity, anxiety-like behavior, spatial working memory, and neuromuscular strength, a series of behavioral tests was performed on all mouse groups during the final week of the intervention. The tests were conducted in the following order, with a 24-hour interval between each test to minimize carry-over effects: Open Field Test, Y-maze Test, and Grip Strength Test. All behavioral procedures were conducted in a dedicated, sound-attenuated behavioral testing room under consistent dim lighting conditions. The mice were allowed to habituate to the testing room for at least 60 minutes before each test session. Trials were recorded and analyzed by the SMART video tracking system (SMART v3.0, Panlab SL, Barcelona, Spain). Grip strength was tested in mouse forelimbs using a grip strength meter (Stoelting Co.).

### Open Field Test

The open field test was employed to assess general locomotor activity and anxiety-like behavior ^42^. The apparatus consisted of a square-shaped white Plexiglas arena (40 cm × 40 cm × 40 cm). Each mouse was gently placed in the center of the arena and allowed to explore freely for 10 minutes. The total distance traveled (in meters) was quantified as an indicator of locomotor activity. The time spent and the distance traveled in the central zone (defined as the central 20 cm × 20 cm area) were measured to evaluate anxiety-like behavior, with decreased activity in the center interpreted as increased anxiety. The arena was thoroughly cleaned with 70% ethanol between trials to eliminate olfactory cues.

### Y-maze Test

Spatial working memory was evaluated using the spontaneous Y-maze test ^43^. The Y-maze was made of black Plexiglas with three identical arms (each 35 cm long × 5 cm wide × 10 cm high) positioned at 120° angles from each other. Each mouse was placed at the end of one designated arm and allowed to explore the maze freely for 10 minutes. An arm entry was recorded when all four paws of the mouse were within the arm. The sequence of arm entries was manually recorded by a blinded experimenter. Spontaneous alternation performance, defined as consecutive entries into all three arms without repetition (e.g., sequence A->B->C), was calculated as a measure of working memory. The percentage of spontaneous alternation was determined using the following formula: Percentage Alternation = [(Number of Alternations) / (Total Arm Entries −2)] × 100%.

### Grip Strength Test

Neuromuscular function was assessed using a grip strength meter ^44^(e.g., Bio-GS3, Bioseb, Vitrolles, France). The forelimb grip strength of each mouse was measured. Briefly, the mouse was held by the tail and allowed to grasp a triangular pull bar connected to the force sensor with its forepaws. The mouse was then pulled backward gently and steadily along the horizontal plane until it released the bar. The peak force (in grams or Newtons) exerted by the mouse before losing its grip was recorded. The test was repeated for five consecutive trials per mouse, with an inter-trial interval of approximately 30 seconds, and the average of the three highest readings was used for statistical analysis.

### Histological analysis of hematoxylin and eosin (H&E) staining

After behavioral testing, euthanize the mice and promptly collect the heart, liver, spleen, lung, kidney, and intestinal tissues. Fix the organization in a 4% paraformaldehyde (PFA) solution at 4 °C for 48 hours. After fixation, the tissue is dehydrated with a series of graded ethanol, removed with xylene, and embedded in paraffin. Subsequently, the embedded tissue coronal sections were sliced into 5 μm thick sections using a rotary slicer (Leica RM2235, Leica Microsystems, Germany). Install the slices on a glass slide coated with poly-L-lysine and dry overnight at 37 ° C. For histological examination, the sections were dewaxed and rehydrated before staining. In short, the slices were deparaffinized twice in xylene for 10 minutes each time, then hydrated for 5 minutes using a decreasing ethanol series, and rinsed in distilled water. Stain with hematoxylin and eosin (H&E) according to the standard protocol, and finally fix with neutral resin. Inspect stained sections under an optical microscope (Olympus BX53, Olympus Corporation, Japan) and capture digital images using the accompanying camera system (DP27, Olympus).

### Quantitative real-time PCR (qPCR) analysis

According to the manufacturer’s instructions, use a commercial RNA isolation kit (TRIzol reagent, Invitrogen, United States) to extract total RNA from frozen brain tissue. Use NanoDrop2000 instruments (ThermoFisher Scientific) to determine the concentration and purity of extracted RNA by spectrophotometry. RNA samples with an A260/A280 ratio between 1.8 and 2.0 and an A260/A230 ratio greater than 2.0 are considered suitable for subsequent experiments. Synthesize the first strand complementary DNA (cDNA) from 1 µg total RNA using a reverse transcription kit (Takara, Japan) in a reaction volume of 20 µL, which includes the step of removing genomic DNA contamination. Perform quantitative PCR using SYBR Green main mixture on a real-time PCR detection system. Each reaction is carried out in a final volume of 20 µL, which contains 10 µL of the main mixture, 0.4 µL of each forward and reverse primer (10 µM), 2 µL of diluted cDNA template, and 7.2 µL of nuclease-free water. The thermal cycling conditions are as follows: initial denaturation at 95 °C for 30 seconds, followed by denaturation at 95 °C for 40 cycles of 5 seconds, and annealing/extension at 60 °C for 30 seconds. Perform melting curve analysis at the end of each run to confirm the specificity of amplification, with a single peak representing a single PCR product. Table 1 lists the sequences of gene-specific primers used in this study.

**Table 1.**
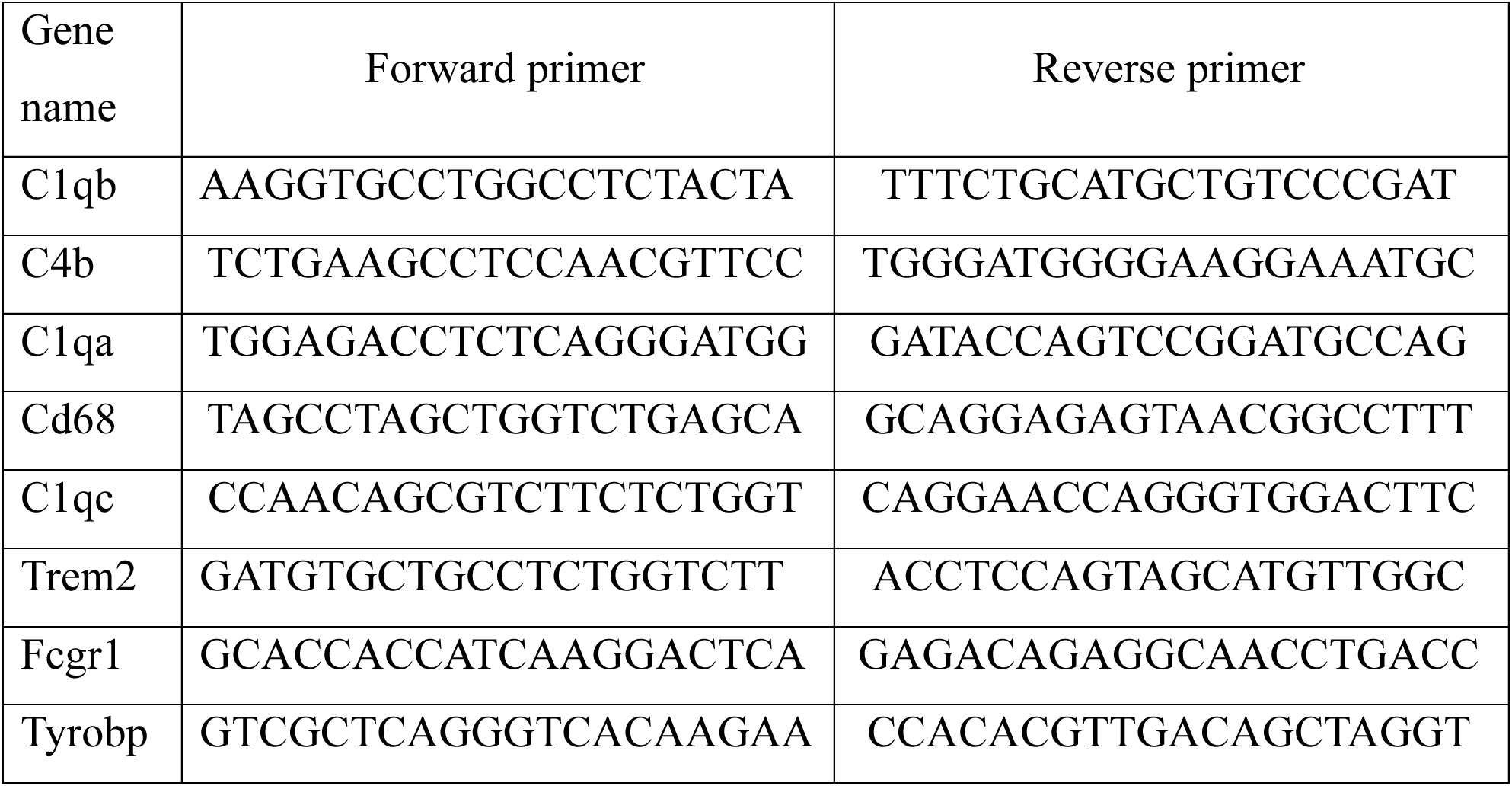
lists the sequences of gene-specific primers.

### Hematological parameters and analysis of liver and kidney function

At the end of the study, blood samples were collected from all mice by drawing blood from the posterior orbital plexus into two types of collection tubes: 1) EDTA-K2 anticoagulant tubes for hematological analysis, and 2) serum separation tubes for biochemical analysis. For complete blood count (CBC), gently invert the whole blood collected in the EDTA-K2 tube several times to ensure proper mixing and analysis within 2 hours after collection. Use an automated hematology analyzer (such as Sysmex XT-2000iV, Sysmex Corporation, Japan) to measure hematological parameters, including white blood cell (WBC) count, red blood cell (RBC) count, hemoglobin (HGB) concentration, hematocrit (HCT), mean corpuscular volume (MCV), mean hemoglobin (MCH), mean corpuscular hemoglobin concentration (MCHC), and platelet (PLT) count. The analyzer is calibrated daily, and quality control is carried out according to the manufacturer’s protocol. To evaluate liver and kidney function, the blood sample in the serum separation tube was coagulated at room temperature for 60 minutes. Subsequently, centrifuge the sample at a speed of 3000 ×g for 15 min at 4 °C to separate the serum. Carefully divide the supernatant (serum) into sterile tubes and store at −80 °C until analysis. Quantify serum levels of alanine aminotransferase (ALT), aspartate aminotransferase (AST), alkaline phosphatase (ACAP), total protein (TP), albumin (ALB), blood urea nitrogen (BUN), and creatinine (CREA) using standard colorimetric or enzyme assay kits (BioVision, USA) on an automated biochemical analyzer (Hitachi 7020, Japan). All analyses were conducted strictly in accordance with the instructions provided by the reagent kit manufacturer.

### Transcriptome sequencing and bioinformatics analysis

Transcriptome sequencing comprises RNA extraction, detection, library construction, and computer sequencing. RNA integrity and concentration were rigorously assessed using an Agilent 2100 Bioanalyzer. Library preparation commenced with the enrichment of polyadenylated mRNA using oligo(dT) beads. The enriched mRNA was fragmented and reverse-transcribed into first-strand cDNA using random hexamer primers and M-MuLV Reverse Transcriptase. Second-strand cDNA was synthesized using DNA Polymerase I and RNase H. The resulting double-stranded cDNA was purified, end-repaired, adenylated, and ligated to Illumina adapters. Fragments of 370–420 bp were size-selected with AMPure XP beads and PCR-amplified. Library quality control included quantification (Qubit 2.0 Fluorometer), size distribution analysis (Agilent 2100 Bioanalyzer), and accurate assessment of effective concentration by qPCR (>2 nM). Finally, libraries were pooled and subjected to 150-bp paired-end sequencing on an Illumina NovaSeq 6000 platform employing the Sequencing-by-Synthesis (SBS) technology. Following sequencing, raw reads were subjected to quality control. The clean reads were then aligned to the reference genome using Hisat2. Gene expression levels were quantified with featureCounts (v1.5.0-p3) to generate read counts for each gene. Based on these counts and gene lengths, FPKM values were calculated to estimate expression abundance. Differential expression analysis between comparative groups was performed using the DESeq2 package (v1.20.0). This method employs a negative binomial generalized linear model to identify statistically significant differences. The resulting p-values were adjusted for multiple testing using the Benjamini-Hochberg procedure to control the false discovery rate (FDR). Functional enrichment analysis was conducted from two complementary perspectives. First, Gene Ontology (GO) enrichment analysis for the differentially expressed genes (DEGs) was carried out using the clusterProfiler package (v3.8.1), with p-values corrected for gene length bias. Terms with an adjusted p-value < 0.05 were considered significantly enriched. Second, Gene Set Enrichment Analysis (GSEA) was implemented using the GSEA desktop tool to evaluate enrichment in pre-defined GO and KEGG gene sets without relying on a fixed DEG threshold. Genes were ranked by their differential expression metric, and significance was determined for gene sets enriched at the top or bottom of the ranked list.

### Proteomics and data analysis

#### Protein extraction

Frozen tissues were pulverized in liquid nitrogen, and the powder was lysed in SDT buffer with 1% DTT. The lysate was vortexed, sonicated on ice, and centrifuged. The supernatant was collected, heated at 95°C, and cooled. Proteins were alkylated with IAM in the dark, precipitated with cold acetone, and the pellet was washed, dried, and redissolved in DB Buffer.

#### FFPE samples

Deparaffinization of FFPE sections was performed using xylene, followed by rehydration through a graded ethanol series and two washes with PBS. After PBS removal, the samples were lysed in protein lysis buffer (4% SDS, 100 mM Tris, pH 7.6) and incubated at 95°C for 10 min with vortexing. Subsequently, the samples were sonicated on ice for 5 min and incubated at 95°C for 60 min for decrosslinking. Proteins were reduced and alkylated by adding TCEP and CAA at 95°C for 5 min. The lysate was centrifuged (12,000 g, 15 min, 4°C), and the supernatant was subjected to acetone precipitation as described in section 2.1.1. The final protein pellet was dissolved in dissolution buffer (6 M urea, 100 mM TEAB, pH 8.5).

#### Protein digestion

An aliquot of each protein sample (up to 100 µL final volume in DB buffer) was digested with trypsin in 100 mM TEAB (pH 8.5) at 37°C for 4 h. The reaction was stopped by acidifying with formic acid (pH < 3). After centrifugation (12,000 g, 5 min, room temperature), the supernatant was desalted using a C18 column. The column was washed three times with a solution of 3% acetonitrile and 0.1% formic acid, and peptides were eluted with 70% acetonitrile containing 0.1% formic acid. The eluate was collected and lyophilized.

#### LC-MS/MS Analysis using Vanquish Neo UHPLC-Astral System

Lyophilized peptides were reconstituted in 10 µL of mobile phase A (0.1% formic acid) and centrifuged (14,000 ×g, 20 min, 4 °C). Subsequently, 200 ng of peptide was injected for analysis. Peptide separation was carried out on a Vanquish Neo UHPLC system using a C18 precolumn (5 mm × 300 µm, 5 µm) and an analytical column (PepMap Neo, 150 µm × 15 cm, 2 µm), both maintained at 50°C. The detailed gradient is provided in Table 1. Mass spectrometry analysis was performed on an Orbitrap Astral instrument equipped with an Easy-spray ion source, operating at a spray voltage of 2.0 kV and an ion transfer tube temperature of 290°C. Data-independent acquisition (DIA) was employed with the following parameters: full MS scans were acquired over m/z 380–980 at a resolution of 240,000, followed by 300 DIA windows (2 Th width) with a normalized collision energy of 25%. MS2 spectra were collected over m/z 150–2000 at a resolution of 80,000, with a maximum injection time of 3 ms. The acquired data were saved in raw format for subsequent analysis.

#### Data analysis

Raw data were processed using DIA-NN software against the *** protein database. Search parameters included: automatic mass tolerance calibration; fixed modification of carbamidomethylation on cysteine; variable modification of N-terminal methionine excision; and a maximum of two missed cleavages. Results were filtered to retain peptides with a Global. Q.Value < 0.01 and proteins with a PG.Q.Value < 0.01. Proteins showing a significant difference (p < 0.05, |log₂FC| > *) between experimental and control groups, as determined by a t-test, were defined as differentially expressed proteins (DEPs).

#### Functional analysis of proteins and DEPs

All identified proteins were functionally annotated using InterProScan for Gene Ontology (GO) and InterPro domains (including Pfam, PRINTS, ProDom, SMART, ProSite, and PANTHER). Protein family and pathway analyses were performed using COG and KEGG databases. For DEPs, volcano plots and hierarchical clustering heatmaps were generated. Functional enrichment analyses were conducted for GO, InterPro, and KEGG pathways. Protein–protein interaction networks were predicted using the STRING database.

### Statistical analysis

Data are expressed as mean ± standard error. Groups were compared using *t*-tests in Prism ver 8.0 (GraphPad Software, Inc., San Diego, CA, USA). Correlation and regression analyses were applied. P < 0.05 defined significance.

## Acknowledgement

This work was supported by Shenzhen Key Laboratory of Magnetic Resonance Physics and Imaging (SYSPG20241211173949066), Shenzhen Medical Academy of Research and Translation (C2301011), and Shenzhen Science and Technology Major Project (KJZD20230923114406014).

---

**Figure S1.**
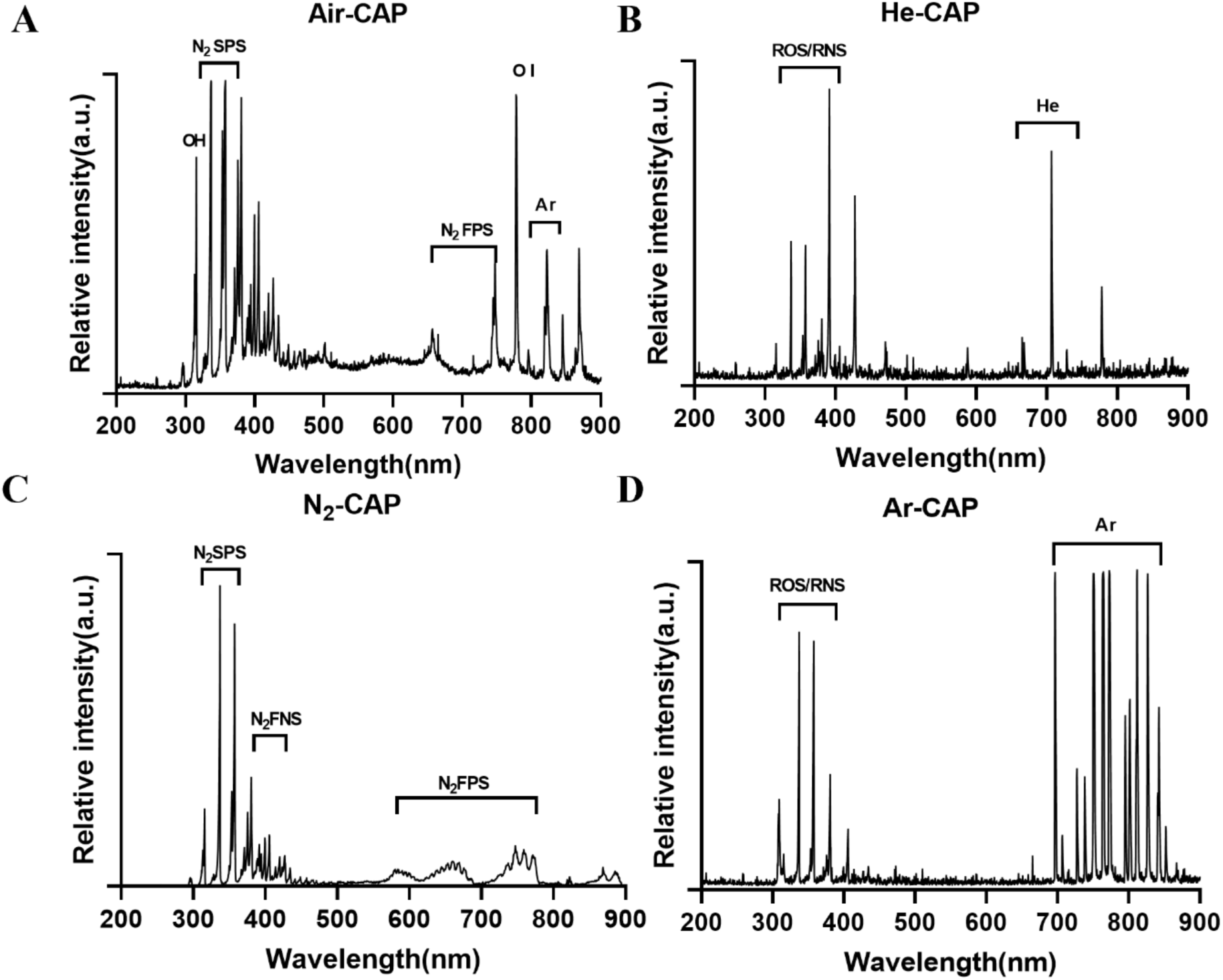
Plasma device diagram and characteristics. (A) Optical emission spectrum(OES) of air source plasma. (B) OES of helium source plasma. (C) OES of nitrogen source plasma. (D) OES of argon source plasma.

**Figure S2.**
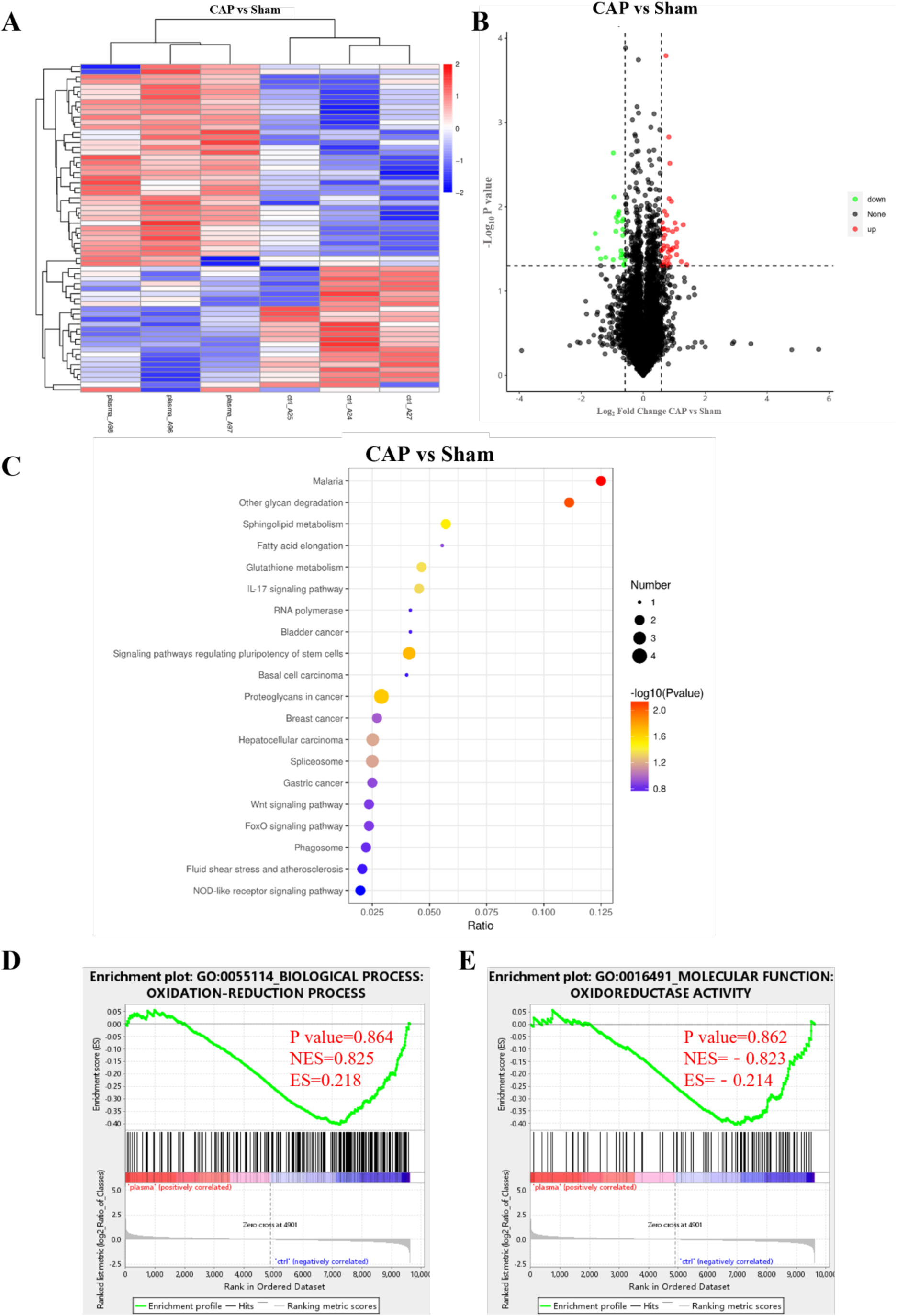
Proteomic analysis of brain proteins in healthy mice treated with CAP. (A) Hierarchical clustering of important proteins. The color scale represents the relative protein level. (B) Volcanic maps of different protein abundances. The proteins that were significantly upregulated (red) and downregulated (blue) were highlighted (| log2FC |>1.5, adjusted p-value<0.05). The dashed line represents the threshold. (C) GO enrichment analysis. Significant enrichment of differentially expressed proteins, ranked by −log10 (adjusted p-value). (D-E) Gene Set Enrichment Analysis (GSEA).

## References

1 Chen, X. et al. Microglia-mediated T cell infiltration drives neurodegeneration in tauopathy. Nature 615, 668–677, doi:10.1038/s41586-023-05788-0 (2023).

2 Graff-Radford, J. et al. New insights into atypical Alzheimer’s disease in the era of biomarkers. Lancet Neurol 20, 222–234, doi:10.1016/S1474-4422(20)30440-3 (2021).

3 Yang, H. et al. Recent Progress in Nanomedicine for the Diagnosis and Treatment of Alzheimer’s Diseases. ACS Nano 18, 33792–33826, doi:10.1021/acsnano.4c11966 (2024).

4 Scheltens, P. et al. Alzheimer’s disease. Lancet 397, 1577–1590, doi:10.1016/S0140-6736(20)32205-4 (2021).

5 Jucker, M. & Walker, L. C. Alzheimer’s disease: From immunotherapy to immunoprevention. Cell 186, 4260–4270, doi:10.1016/j.cell.2023.08.021 (2023).

6 Self, W. K. & Holtzman, D. M. Emerging diagnostics and therapeutics for Alzheimer disease. Nat Med 29, 2187–2199, doi:10.1038/s41591-023-02505-2 (2023).

7 Liu, N., Liang, X., Chen, Y. & Xie, L. Recent trends in treatment strategies for Alzheimer(’)s disease and the challenges: A topical advancement. Ageing Res Rev 94, 102199, doi:10.1016/j.arr.2024.102199 (2024).

8 Park, M. et al. Effects of transcranial ultrasound stimulation pulsed at 40 Hz on Abeta plaques and brain rhythms in 5xFAD mice. Transl Neurodegener 10, 48, doi:10.1186/s40035-021-00274-x (2021).

9 Murdock, M. H. et al. Multisensory gamma stimulation promotes glymphatic clearance of amyloid. Nature 627, 149–156, doi:10.1038/s41586-024-07132-6 (2024).

10 Iaccarino, H. F. et al. Gamma frequency entrainment attenuates amyloid load and modifies microglia. Nature 540, 230–235, doi:10.1038/nature20587 (2016).

11 Martorell, A. J. et al. Multi-sensory Gamma Stimulation Ameliorates Alzheimer’s-Associated Pathology and Improves Cognition. Cell 177, doi:10.1016/j.cell.2019.02.014 (2019).

12 Hess, B. L., Piazolo, S. & Harvey, J. Lightning strikes as a major facilitator of prebiotic phosphorus reduction on early Earth. Nat Commun 12, 1535, doi:10.1038/s41467-021-21849-2 (2021).

13 Brune, W. H. et al. Extreme oxidant amounts produced by lightning in storm clouds. Science 372, 711–715, doi:10.1126/science.abg0492 (2021).

14 Liao, P., Kang, J., Xiang, R., Wang, S. & Li, G. Electrocatalytic Systems for NO(x) Valorization in Organonitrogen Synthesis. Angew Chem Int Ed Engl 63, e202311752, doi:10.1002/anie.202311752 (2024).

15 Zhang, X. et al. Spaceborne Observations of Lightning NO(2) in the Arctic. Environ Sci Technol 57, 2322–2332, doi:10.1021/acs.est.2c07988 (2023).

16 Jiang, H. J. et al. Mimicking lightning-induced electrochemistry on the early Earth. Proc Natl Acad Sci U S A 121, e2400819121, doi:10.1073/pnas.2400819121 (2024).

17 Jenewein, C., Maíz-Sicilia, A., Rull, F., González-Souto, L. & García-Ruiz, J. M. Concomitant formation of protocells and prebiotic compounds under a plausible early Earth atmosphere. Proc Natl Acad Sci U S A 122, e2413816122, doi:10.1073/pnas.2413816122 (2025).

18 Zhao, Y., Guo, Q., Tian, J., Liu, W. & Wang, X. TREM2 bridges microglia and extracellular microenvironment: Mechanistic landscape and therapeutical prospects on Alzheimer’s disease. Ageing Research Reviews 103, 102596, doi:10.1016/j.arr.2024.102596 (2025).

19 Kodali, M. et al. Residual microglia following short-term PLX5622 treatment in 5xFAD mice exhibit diminished NLRP3 inflammasome and mTOR signaling, and enhanced autophagy. Aging Cell 24, e14398, doi:10.1111/acel.14398 (2024).

20 Qiao, L. et al. H2O2-responsive multifunctional nanocomposite for the inhibition of amyloid-β and Tau aggregation in Alzheimer’s disease. BMEMat 1, e12011, 10.1002/bmm2.12011 (2023).

21 Li, Y. et al. 3D-cultured BMSC exosomes improve cerebral ischemia/reperfusion injury-induced neuronal apoptosis by regulating the microglia polarization. BMEMat 3, e70000, 10.1002/bmm2.70000 (2025).

22 Wang, S. et al. TREM2 drives microglia response to amyloid-β via SYK-dependent and -independent pathways. Cell 185, doi:10.1016/j.cell.2022.09.033 (2022).

23 Fu, D. et al. Cold atmospheric plasma as a novel “drug” for cancer therapy. J Control Release 386, 114118, doi:10.1016/j.jconrel.2025.114118 (2025).

24 Fang, T., Chen, Z. & Chen, G. Advances in cold atmospheric plasma therapy for cancer. Bioact Mater 53, 433–458, doi:10.1016/j.bioactmat.2025.07.031 (2025).

25 Shinzato, G. T. et al. Non-invasive sound wave brain stimulation with Transcranial Pulse Stimulation (TPS) improves neuropsychiatric symptoms in Alzheimer’s disease. Brain Stimulation 17, 413–415, doi:10.1016/j.brs.2024.03.007 (2024).

26 Chen, G. et al. Portable air-fed cold atmospheric plasma device for postsurgical cancer treatment. Science Advances 7, eabg5686, doi:10.1126/sciadv.abg5686 (2021).

27 Fang, T., Cao, X., Shen, B., Chen, Z. & Chen, G. Injectable cold atmospheric plasma-activated immunotherapeutic hydrogel for enhanced cancer treatment. Biomaterials 300, 122189, doi:10.1016/j.biomaterials.2023.122189 (2023).

28 Chen, Z. T., Obenchain, R. & Wirz, R. E. Tiny Cold Atmospheric Plasma Jet for Biomedical Applications. Processes 9, doi:10.3390/pr9020249 (2021).

29 Li, Q. et al. Impaired lipophagy induced-microglial lipid droplets accumulation contributes to the buildup of TREM1 in diabetes-associated cognitive impairment. Autophagy 19, 2639–2656, doi:10.1080/15548627.2023.2213984 (2023).

30 Li, J. et al. Combination of autophagy and NFE2L2/NRF2 activation as a treatment approach for neuropathic pain. Autophagy 17, 4062–4082, doi:10.1080/15548627.2021.1900498 (2021).

31 Li, F. et al. Low-intensity pulsed ultrasound stimulation (LIPUS) modulates microglial activation following intracortical microelectrode implantation. Nat Commun 15, 5512, doi:10.1038/s41467-024-49709-9 (2024).

32 Prichard, A. et al. Brain rhythms control microglial response and cytokine expression via NF-kappaB signaling. Sci Adv 9, eadf5672, doi:10.1126/sciadv.adf5672 (2023).

33 Tao, L. et al. Microglia modulation with 1070-nm light attenuates Abeta burden and cognitive impairment in Alzheimer’s disease mouse model. Light Sci Appl 10, 179, doi:10.1038/s41377-021-00617-3 (2021).

34 Li, Y. et al. Ultrasound Controlled Anti-Inflammatory Polarization of Platelet Decorated Microglia for Targeted Ischemic Stroke Therapy. Angew Chem Int Ed Engl 60, 5083–5090, doi:10.1002/anie.202010391 (2021).

35 Lin, Y. et al. Electromagnetic pulse exposure induces neuroinflammation and blood-brain barrier disruption by activating the NLRP3 inflammasome/NF-kappaB signaling pathway in mice. Ecotoxicol Environ Saf 292, 117972, doi:10.1016/j.ecoenv.2025.117972 (2025).

36 Chen, G. et al. Transdermal cold atmospheric plasma-mediated immune checkpoint blockade therapy. Proceedings of the National Academy of Sciences of the United States of America 117, 3687–3692, doi:10.1073/pnas.1917891117 (2020).

37 Cao, X. et al. RSL3-loaded nanoparticles amplify the therapeutic potential of cold atmospheric plasma. J Nanobiotechnology 23, 136, doi:10.1186/s12951-025-03211-6 (2025).

38 Liu, Y. et al. Matrix stiffness-dependent microglia activation in response to inflammatory cues: in situ investigation by scanning electrochemical microscopy. Chem Sci 15, 171–184, doi:10.1039/d3sc03504b (2023).

39 Chen, C. et al. Exosomes Derived from M2 Microglial Cells Modulated by 1070-nm Light Improve Cognition in an Alzheimer’s Disease Mouse Model. Adv Sci (Weinh) 10, e2304025, doi:10.1002/advs.202304025 (2023).

40 Lv, Z. et al. Clearance of beta-amyloid and synapses by the optogenetic depolarization of microglia is complement selective. Neuron 112, 740–754.e747, doi:10.1016/j.neuron.2023.12.003 (2024).

41 Zhao, Y. et al. TREM2 Is a Receptor for β-Amyloid that Mediates Microglial Function. Neuron 97, doi:10.1016/j.neuron.2018.01.031 (2018).

42 Qiu, S. et al. Adult-onset CNS myelin sulfatide deficiency is sufficient to cause Alzheimer’s disease-like neuroinflammation and cognitive impairment. Molecular Neurodegeneration 16, 64, doi:10.1186/s13024-021-00488-7 (2021).

43 Lv, S. et al. Integrated dopamine sensing and 40 Hz hippocampal stimulation improves cognitive performance in Alzheimer’s mouse models. Nature Communications 16, 5948, doi:10.1038/s41467-025-60903-1 (2025).

44 Bellantuono, I. et al. A toolbox for the longitudinal assessment of healthspan in aging mice. Nat Protoc 15, 540–574, doi:10.1038/s41596-019-0256-1 (2020).

